# High throughput sequencing of multiple amplicons for barcoding and integrative taxonomy

**DOI:** 10.1101/073304

**Authors:** Perrine Cruaud, Jean-Yves Rasplus, Lillian Jennifer Rodriguez, Astrid Cruaud

## Abstract

Until now, the potential of NGS has been seldom realised for the construction of barcode reference libraries. Using a two-step PCR approach and MiSeq sequencing, we tested a cost-effective method and developed a custom workflow to simultaneously sequence multiple markers (*COI, Cytb* and *EF*, altogether 2kb) from hundreds of specimens. Interestingly, primers and PCR conditions used for Sanger sequencing did not require optimisation to construct MiSeq library. After completion of quality controls, 87% of the species and 76% of the specimens had valid sequences for the three markers. Nine specimens (3%) exhibited two divergent (up to 10%) sequence clusters. In 95% of the species, MiSeq and Sanger sequences obtained from the same samplings were similar. For the remaining 5%, species were paraphyletic or the sequences clustered into two divergent groups (>7%) on the final trees (Sanger + MiSeq). These problematic cases are difficult to explain but may represent coding NUMTS or heteroplasms. These results highlight the importance of performing quality control steps, working with expert taxonomists and using more than one marker for DNA-taxonomy or species diversity assessment. The power and simplicity of this method appears promising to build on existing experience, tools and resources while taking advantage of NGS.

## INTRODUCTION

While next-generation sequencing (NGS) is commonly used to analyse bulk environmental samples (metabarcoding) ^1-3^, Sanger sequencing remains the standard approach in generating DNA barcode libraries ^4^. This is unfortunate as the cost-effective acquisition of barcode sequences from hundreds of specimens identified to species by expert taxonomists could accelerate the construction of accurate reference libraries and increase their completeness^2,5^.

As it generates up to 25 million paired end reads (2*300bp), the Illumina MiSeq platform makes possible the sequencing of several hundreds of individuals on a set of informative barcodes. This allows for the increase not only in the number of species but also in the number of specimens included in reference databases, which is crucial, as a better coverage of the geographical range of the species and a better characterisation of the intraspecific variability lead to more accurate identification^6,7^. Two-step polymerase chain reactions (PCR) are easy methods that can be used to generate amplicon libraries for Illumina sequencing. In the first PCR reaction the targeted DNA region is amplified using specific primers flanked by tails (Fig. 1). These tails allow for a second PCR reaction to add Illumina adaptor sequences and indexes to multiplex samples ^8^. Theoretically, two-step PCR approaches provide an opportunity to build on existing experience and tools (e.g. primers and PCR conditions), which make them very attractive.

**Figure 1.**
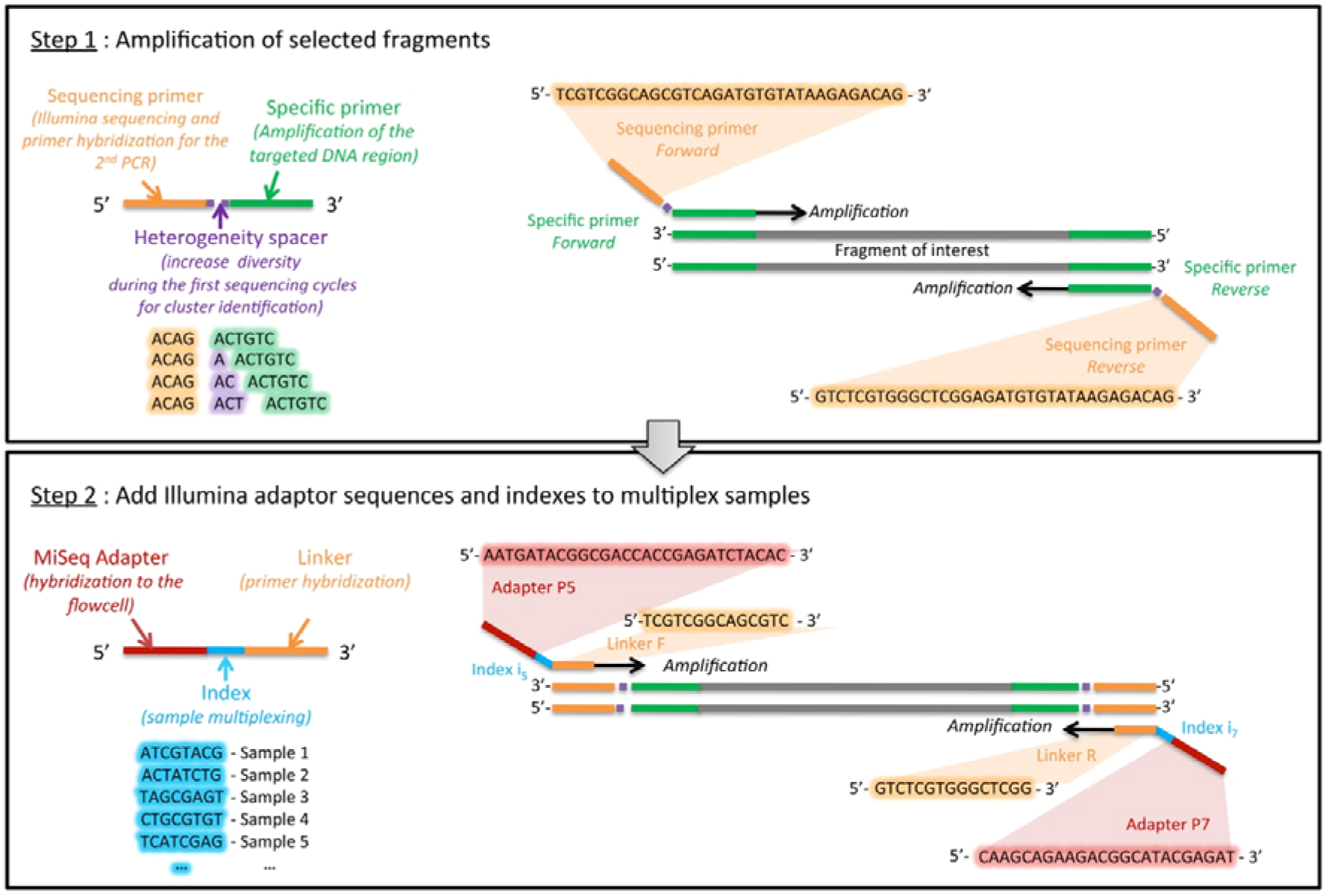
Illustration of the two-step PCR approach.

Combining two-step PCR approaches and high-throughput sequencing may contribute to circumvent some of the main pitfalls of barcoding revealed by many studies ^9^. Indeed, heteroplasms ^10,11^; NUMTS (NUclear MiTochondrial DNA segments) ^12^, endosymbionts ^13^, parasitoids 14 or contaminants may be sometimes preferentially amplified by the primer pair used and are frequently sequenced using Sanger methods. Using NGS, these non-target loci may be simultaneously amplified with the targeted COI, sequenced within the sequencing depth and better identified by post sequencing analyses. Furthermore, combining two-step PCR and MiSeq sequencing may also help to increase the number of genes sequenced for barcoding. Indeed to circumvent the main pitfalls associated with the use of a single, mitochondrial gene, it has been acknowledged that an increase in the number of genes analysed is desired, though most studies still rely on *COI* only^9^. This increase is even more recommended when it comes to DNA-based species delimitation ^15-17^ or phylogeography. However, the addition of loci often comes at the expense of sampling ^18^. By combining multiplexing techniques with high throughput sequencing, researchers may no longer need to choose between more samples or more characters. Finally, adding one or a few nuclear genes aside the standard mitochondrial fragment (*COI*) may facilitate the identification of mtDNA introgression ^19^.

Recently, genome skimming, the low-coverage shotgun sequencing of total genomic DNA ^20^ has been proposed as a next generation barcoding tool ^21^. However, a switch to databases including the complete genome sequence of each organism on Earth is still unrealistic due to unaffordable costs. Furthermore high consumable costs, increased demands on data storage, analytical issues, as well as potential difficulties in obtaining material transfer agreements ^21^, challenge the implementation of this method. In any case, identification of random scaffolds is not possible with current databases. Thus, when genome skimming was used to capture the genomic diversity of bulk arthropod samples ^22^, *ca* 70% of the recovered scaffolds could not be identified to species with existing databases. Therefore, there should be a gradual and step-wise implementation of genome skimming.

In this light, taking advantage of current databases seems more realistic, especially to make use of the huge effort undertaken over the past 15 years in compiling millions of *COI* sequences for hundreds of thousands species (e.g. the International Barcode of Life project iBOL). Finally, in many groups of living organisms, *COI* or a couple of genetic markers provide an accurate identification, even if problems do exist in some groups ^23-25^.

Here, we focused on a group of chalcid wasps (Hymenoptera, Agaonidae, *Ceratosolen*) for which we have accumulated Sanger sequences on two mitochondrial [*COI* and cytochrome b (*Cytb*)], and one nuclear markers [elongation factor-1a (*EF1a*)], over the past 20 years and on which we have a strong taxonomic expertise that is essential to detect mismatches between morphological and molecular identification. Using a two-step PCR approach (Fig. 1) and Illumina MiSeq sequencing, we amplified and sequenced the same three markers (Table 1) on 115 species of *Ceratosolen* (369 specimens). We process raw data using a custom workflow including quality control steps (Figs 2&3) and compared our results to the Sanger data set.

**Figure 2.**
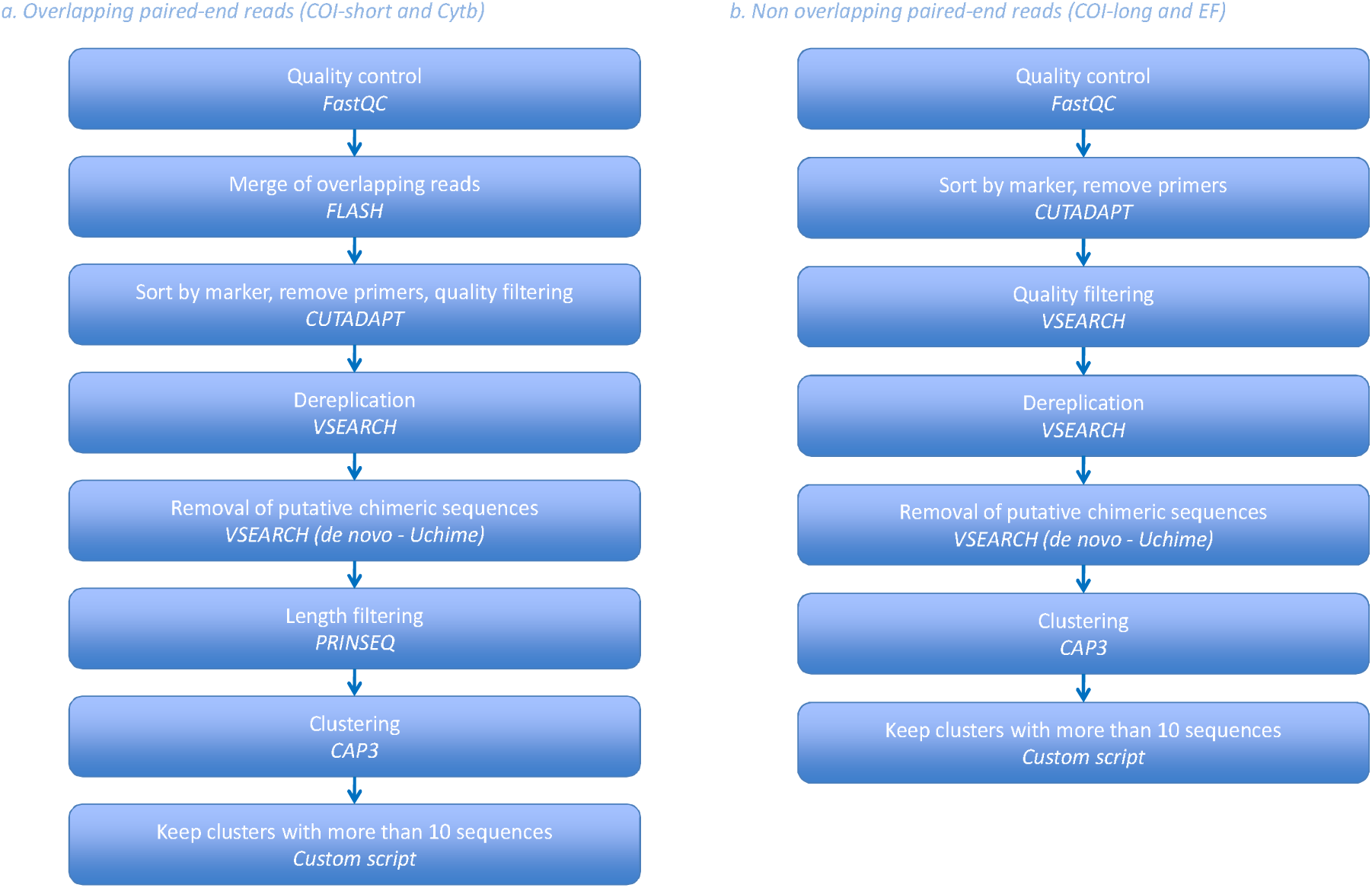
Analytical workflow. Step 1, from read filtering to clustering.

**Figure 3.**
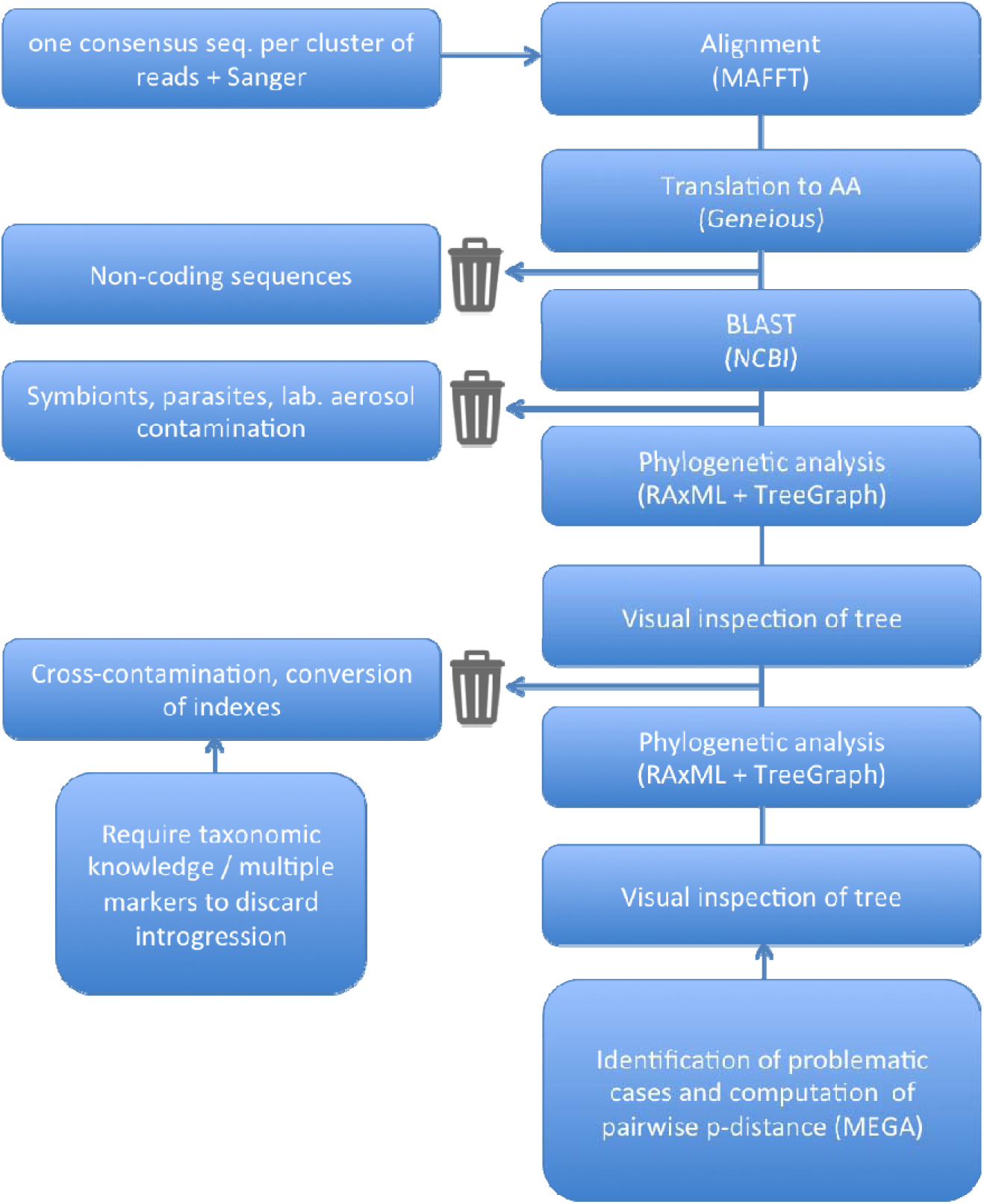
Analytical workflow. Step 2, quality control of clusters of reads.

**Table 1.**
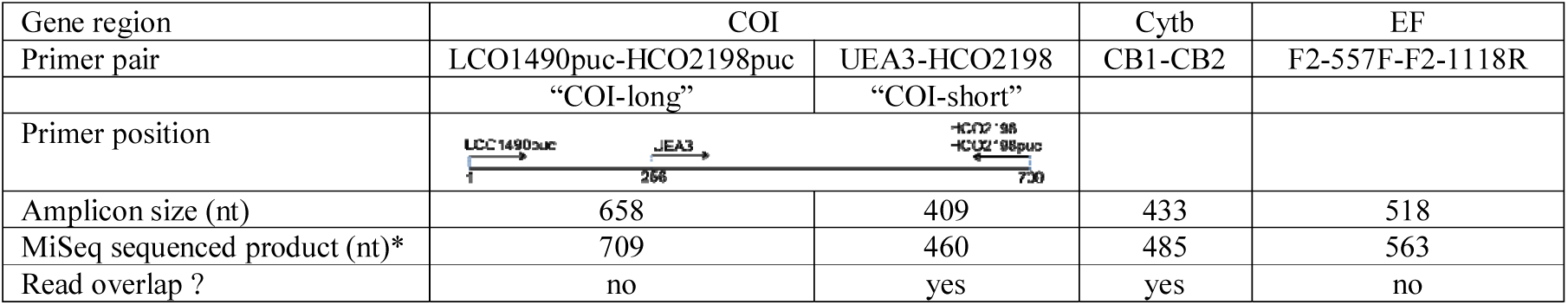
DNA regions targeted for amplification. (* sequenced product = forward primer + amplicon + reverse primer)

The first objective of this study was to test the feasibility of the method. Then, we wanted to determine the best strategy to analyse MiSeq raw data for reference database construction and/or DNA-based identification. Indeed, with the thousands of sequences per sample produced by the MiSeq platform, sequence correction is not a burden anymore, but other issues may appear that need to be considered. On the one hand, thank to sequencing depth, chances of actually getting sequences of the target locus are higher compared to Sanger sequencing. On the other hand, non-target loci (i.e. pseudogenes, heteroplasmic sequences) are also sequenced and target DNA region must be sorted out from the rest of the sequences. More specifically, one may wonder whether the cluster that contains the largest proportion of reads always corresponds to the targeted loci. Two studies suggested it might be so in most cases ^2,4^, but other analyses are required. Finally, at some point, Sanger and Illumina sequences will both be used in reference databases, for integrative taxonomy, or for DNA-based identification of specimen. Consequently, identifying potential issues during data reconciliation was the third objective of this study.

## RESULTS

### MiSeq library construction and sequencing

Amplification success for each gene region (bands on the gel at the expected size after the first PCR step) is summarized in Tables 2&3. The success of PCR was higher for the mitochondrial genes. A PCR amplification product was observed for 80.9% of the species for *COI*-long, 86.1% for *COI*-short, 85.2 % for *Cytb*, and 77.4% for *EF*. As might be expected, we found a negative correlation between amplification success and time elapsed since specimen collection (Fig. 4). Interestingly, the overall amplification success between Sanger and MiSeq data sets were similar, though longer primers were used in the two-step PCR approach. DNA extraction seemed to have failed for 47 specimens (no PCR amplification product visible on gel). Analyses of the per-sequence quality scores showed that the sequencing quality of 40,1% (resp. 25.9%) of the forward (resp. reverse) reads reached Q30. We observed increased error rates towards the end of the reads (especially reverse reads). As a consequence, the paired reads did not overlap for *EF*, though the sequenced product (563bp) fall into the range of the MiSeq Reagent Kits v3. A total of 18,688,278 Illumina paired-end reads were obtained with an average number of raw reads per sample of 38,913 (range = 673 - 158,278).

**Figure 4.**
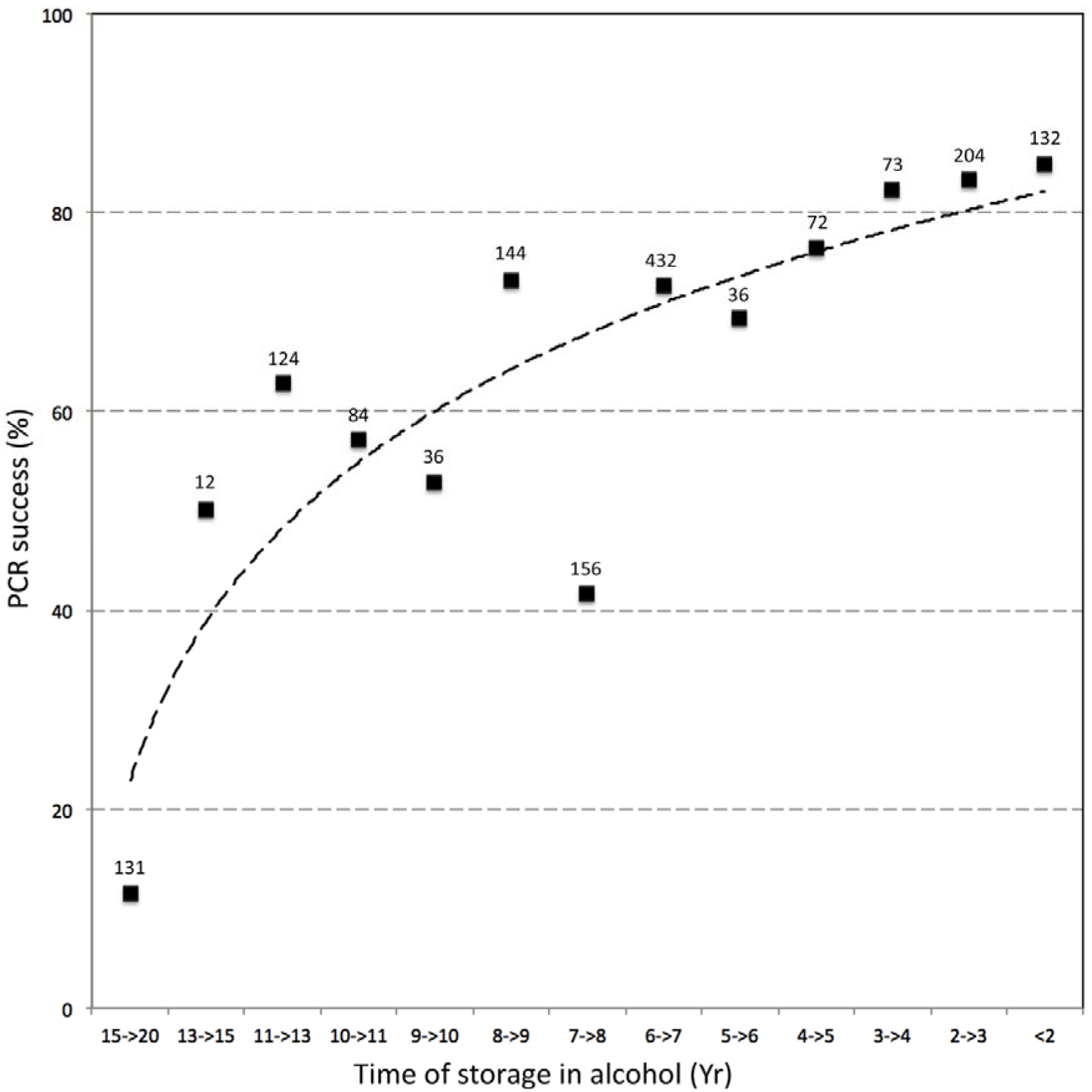
Success rate of amplification as a factor of time since storage of specimens in alcohol.

**Table 2.** Sequencing results of the MiSeq data set. (Regions for which paired-end reads did overlap)

| Gene region | COI-short | Cytb |
| --- | --- | --- |
| PCR success (MiSeq)<sup>1</sup> | 272 (73.7%)<br>99 species (86.1%) | 272 specimens (73.7%)<br>98 species (85.2%) |
| Nb (%) of specimens for which at least one cluster of reads was obtained | 325 (88.1%)<br>[incl. 58 with PCR-] | 280(75.9%)<br>[incl. 17 with PCR-] |
| Nb of specimens with PCR+ but no cluster of reads | 5 (1.8%) | 9 (3.3%) |
| Average (maximum) nb of clusters per specimen | 2.2 (12) | 1.6 (8) |
| Nb of specimens for which the consensus sequence of at least one cluster successfully passed the translation to AA step | 270 (83.1%)<br>[incl. 39 with PCR-] | 270 (96.4%)<br>[incl. 15 with PCR-] |
| Nb of specimens for which the consensus sequence of the first major cluster<sup>2</sup> did not pass the translation to AA step | 77 (23.7%)<br>[incl. 25 with PCR -] | 17 (6.1%)<br>[incl. 2 with PCR -] |
| Nb of specimens for which the consensus sequence of the major cluster<sup>3</sup> did not belong to the target group<sup>4</sup> | 0 | 1 (0.4%)<br>[incl. 1 with PCR -]<br>(lab. aerosol contamination) |
| Nb of specimens for which the consensus sequence of the major cluster was identical to sequence(s) of another species of *Ceratosolen*<sup>5</sup> | 21 (6.5 %)<br>[incl. 9 with PCR -] | 8 (2.9%)<br>[incl. 1 with PCR -] |

<sup>1</sup>as revealed by a visual inspection of the gel after the first PCR step
<sup>2</sup>The cluster that contains the largest proportion of reads/sequences is called the “major cluster”.
<sup>3</sup>At this stage of the process, “major cluster” stands for the cluster that contains the largest proportion of reads/sequences AND whose consensus sequence successfully passed the translation to amino acids step.
<sup>4</sup>as revealed by NCBI-BLAST (e.g. symbionts, parasites, predators or laboratory aerosol contamination)
<sup>5</sup>as revealed by visual inspection of trees. In this case, sequences belong to the target group (*Ceratosolen*) but are identical to sequences from another species (100% BP). May be due to cross-contamination during library preparation or conversion of indexes due to mixed clusters on the flow cell (clonal clusters derived from more than one template molecule), but also to mtDNA introgression (which is undetectable without taxonomic knowledge of the group or comparison with nuDNA data sets).

**Table 3.** Sequencing results of the MiSeq data set. (Regions for which paired-end reads did not overlap). See Table 2 for legends Gene region.

| Gene region | COI-long |  | EF |  |
| --- | --- | --- | --- | --- |
|  | Forward reads | Reverse reads | Forward reads | Reverse reads |
| PCR success (MiSeq)<sup>1</sup> | 244 specimens (66.1%)<br>93 species (80.9%) |  | 244 specimens (66.1%)<br>89 species (77.4%) |  |
| Nb (%) of specimens for which at least one cluster of reads was obtained | 273 (74.0%)<br>[incl. 36 with PCR-] | 305 (82.7%)<br>[incl. 72 with PCR-] | 300 (81.3%)<br>[incl. 61 with PCR-] | 330 (89.4%)<br>[incl. 93 with PCR-] |
| Nb of specimens with PCR+ but no cluster of reads | 7 (2.9%) | 11 (4.5%) | 5 (2.0%) | 7 (2.9%) |
| Average (maximum) nb of clusters per specimen | 3,0 (17) | 3,5 (12) | 3,7 (17) | 4,3 (23) |
| Nb of specimens for which the consensus sequence of at least one cluster successfully passed the translation to AA step | 218 (79.9%)<br>[incl. 18 with PCR-] | 199 (65.2%)<br>[incl. 25 with PCR-] | 291 (97.0%)<br>[incl. 50 with PCR-] | 227 (68.8%)<br>[incl. 27 with PCR-] |
| Nb of specimens for which the consensus sequence of the first cluster<sup>2</sup> did not pass the translation to AA step | 116 (42.5%)<br>[incl. 25 with PCR-] | 278 (91.1%)<br>[incl. 63 with PCR-] | 33 (11.0%)<br>[incl. 20 with PCR-] | 279 (84.5%)<br>[incl. 86 with PCR-] |
| Nb of specimens for which the consensus sequence of the major cluster<sup>3</sup> did not belong to the target group<sup>4</sup> | 28(10.3%)<br>[incl. 4 with PCR-]<br>(27 nematods, 1 lab. aerosol contamination) | 47 (15.4%)<br>[incl. 2 with PCR-]<br>(41 *Wolbachia*, 6 nematods) | 0 | 0 |
| Nb of specimens for which the consensus sequence of the major cluster was identical to sequence(s) of another species of *Ceratosolen*<sup>5</sup> | 6 (2.2%)<br>[incl 2 with PCR-] | 1 (0.3%)<br>[incl 0 with PCR-] | 11 (3.7%)<br>[incl. 4 with PCR-] | 6 (1.8%)<br>[incl. 2 with PCR-] |

### Quality control of clusters of reads

The number of clusters of reads varied among samples and genes (Tables 2 & 3). More clusters were obtained when paired-end reads did not overlap, probably because of the increased error rates towards the end of the reads. After completion of our workflow (Figs 2&3; Table 4), ca 76% of the specimens have a sequence for the three-targeted genes and at least one sequence was retained for 94.8% (*COI*), 82.6% (*Cytb*) and 84.3% (*EF*) of the species. Sequencing what appeared as negative PCR amplifications allowed saving up to 11 species for *EF*, 9 for *COI* and 3 for *Cytb*. On the other hand, no sequence were obtained for about 6.6% (*Cytb*), 5.1 % (*COI*) and 3.7% (*EF*) of the samples for which an amplicon was visible on the gel (Table 4).

**Table 4.**
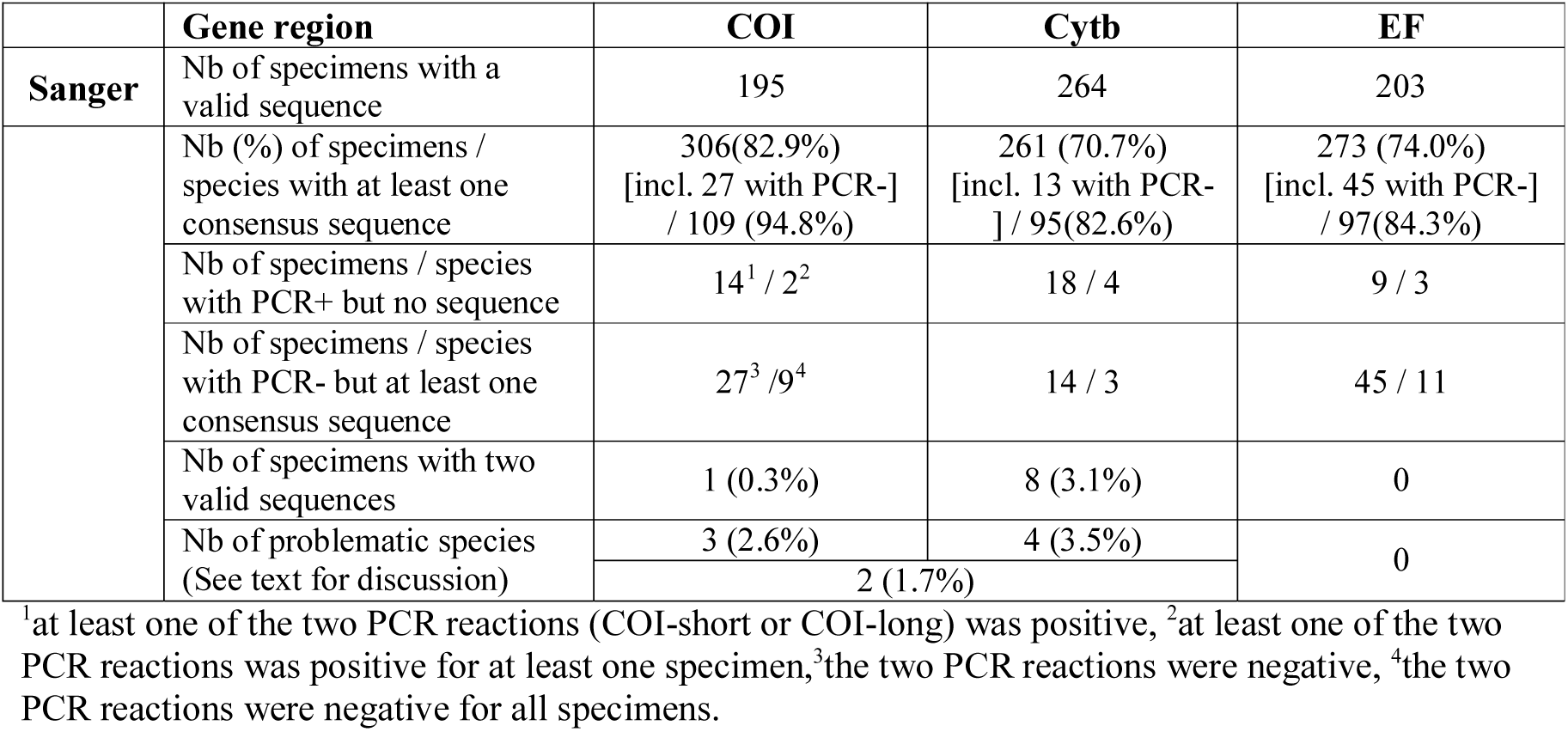
Final results obtained on the combined data set (Sanger + MiSeq), after completion of the workflow.

Translation to amino acids showed that 57.2% (*COI*-long, forward reads), 8.9% (*COI*-long, reverse reads), 76.3% (*COI*-short), 93.9% (*Cytb*), 89.0% (*EF*, forward reads), 15.5% (*EF*, reverse reads) of the major clusters obtained from positive PCR were coding (Tables 2 & 3). Among clusters obtained from positive PCR and that passed the translation step, 100% of the *COI*-short, *Cytb*, *EF* clusters as well as 89.7% (*COI*-long, forward reads) and 84.6% (*COI*-long, reverse reads) clusters blasted with Agaonidae sequences on NCBI. Non-homolog sequences mostly belong to symbionts (*Wolbachia*) or parasites (nematodes). Finally, among clusters obtained from positive PCR products and that passed the translation step, an average of 2.6% only had a consensus sequence identical to another species of *Ceratosolen* and may represent contamination or conversion of indexes. Therefore, the cluster that contained the largest proportion of reads did not necessarily represent a valid sequence.

After completion of the workflow, a few specimens (2.4%) were represented by two consensus sequences in the final MiSeq data set: one specimen for *COI*, for which sequences of *COI*-long and *COI*-short were different (JRAS03502_0153, Fig. S1) and eight specimens for *Cytb*, for which the major and the second major clusters had different consensus sequences (JRAS02196_0155, 56; JRAS01683_0151, 55, 56; JRAS02370_0151, 55, 56; Fig. S2). Phylogenetic inference revealed that one of the copies was (almost) identical to Sanger sequences while the other clustered apart with an average pairwise sequence divergence ranging from 7.3% to 10.3% (Fig. 4, S1, S2, Table 4). These cases are problematic as no objective criteria allow the removal of one of the sequences from the final data set. Lastly, when combining Sanger and MiSeq datasets, 9 species formed paraphyletic assemblages or clustered into two divergent (>7%) groups of sequences: 3 on the *COI* tree (Fig. S1), 4 on the *Cytb* tree (Figs 5, S2), 2 on both *COI* and *Cytb* trees, Table 4). Although two copies of *EF* have been reported in Hymenoptera ^26^, no problematic case was detected on the tree obtained from the analysis of the *EF* data set (Fig. S3, Table 4).

**Figure 5.**
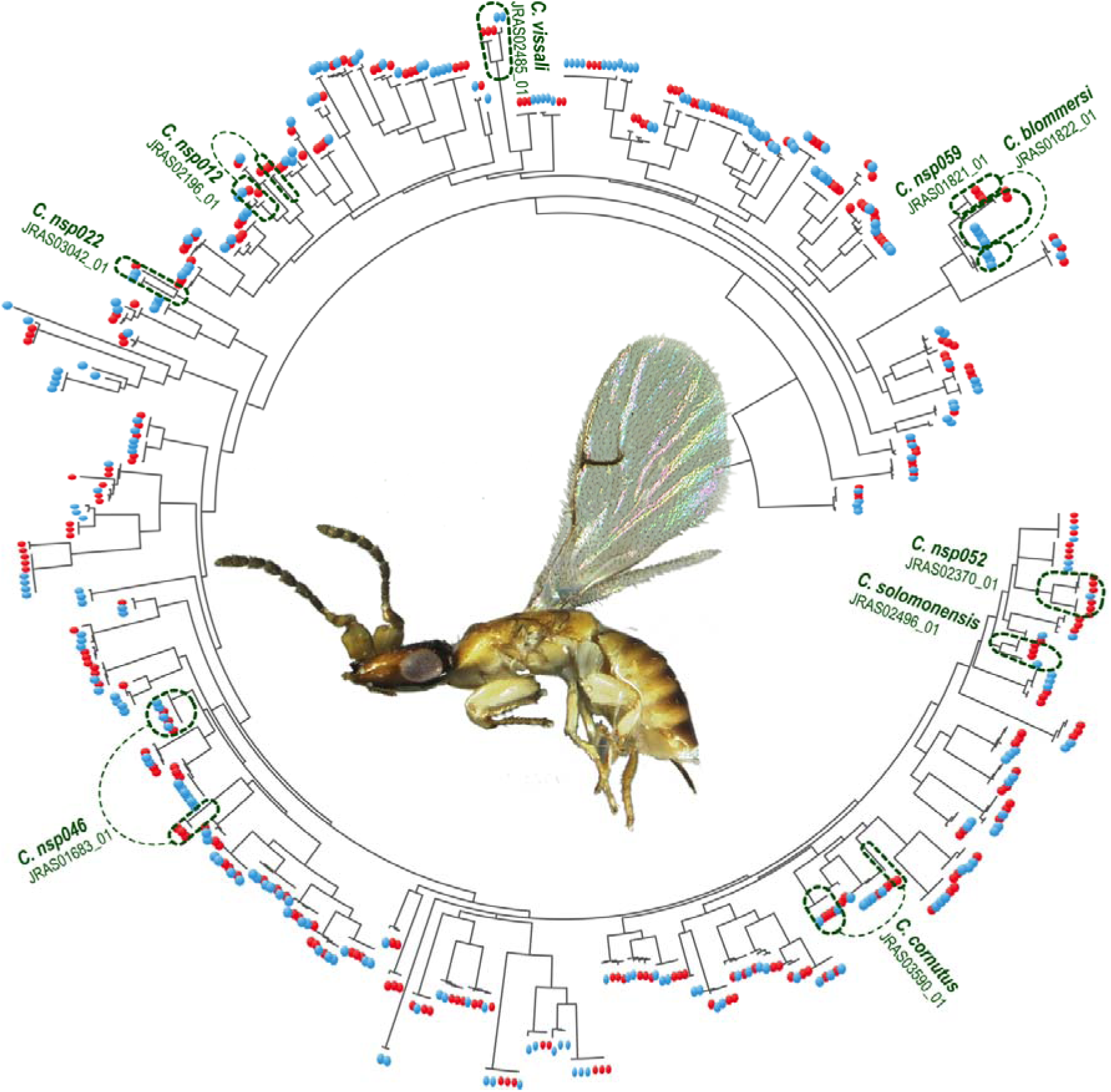
RAxML tree for the Cytb data set (MiSeq+Sanger) (BP : 1000 replicates). Red (resp. blue) circles represent sequences produced by Sanger (resp. MiSeq) sequencing. Dotted lines show problematic cases as discussed in text (see also Fig S2).

## DISCUSSION

In this study, we showed that a two-step PCR approach followed by Illumina sequencing may help to increase the number of sampled species and specimens in existing barcode databases. By including nuclear genes, this method could allow accurate identification of specimens within species complexes where mitochondrial markers may be misleading (e.g. introgression^6,7^). Interestingly, primers and PCR conditions used to generate Sanger data sets did not need optimisation in order to be used for MiSeq library preparation. Moreover, this approach does not require costly investments in laboratory equipment and supplies. Provided that adapters/index and primers are compatible (e.g. no hairpin structure), researchers can keep on working with markers they have previously selected for informativeness^27^.

Increased error rates towards the end of the reads (especially reverse reads) have made the bioinformatic processing of data less convenient with a necessary switch to algorithms that allow clustering of sequences with different length. Nevertheless, processing remains feasible and fast with available programs (48 hours were required on 8-cores of a 16-cores Linux, 2.9GHz, 64GB RAM computer to process raw data). This increase in error rate could be due to accumulation of phasing and pre-phasing events throughout the sequencing process ^28^. When contacted, the Illumina technical support team acknowledged the issue and informed us that they were working to fix it. Progress is being made and work associated with resolving this issue is continuing. It would thus be reasonable to think that in the next couple of years read length will increase and amplicons of any size could be sequenced.

Results from this pilot study suggest that it may be helpful to sequence PCR products with no visible bands on the gel (potentially negative PCR). Indeed, we obtained sequences that passed our quality control steps for 86 specimens for which no amplification product was detected on the gel. When using Sanger method, sequencing what seemed to be negative PCR products was discouraged because of sequencing cost. This aspect now becomes affordable with NGS methods. From a practical point of view, pooling positive and negative PCR products can lead to low concentration of the library (<2nM before denaturation, which is the minimum concentration recommended by the Illumina protocol). Here we obtained 0.24 nM but used Tris-HCL to neutralize the increased NaOH concentration as suggested in the NextSeq protocol of Illumina. Neutralization should also work for other MiSeq libraries provided than PCR success is not too low (here the average PCR success was 70%). As for the classical PCR approach, the negative correlation observed between amplification success and time elapsed since specimen collection argues in favour of rapid DNA extraction after fieldwork instead of long-term storage of specimens in EtOH.

In this study, we amplified three genomic regions on 369 samples but other experimental design may be used to better fit with researcher needs (more markers with less samples or more samples with less markers). The ca 25M reads generated by the MiSeq platform indeed open up many possibilities. That says, sequencing depth allocated to each marker should be large enough to allow sequencing of the target region. Indeed, sequencing depth of NGS methods statistically alleviates the effect of base calling errors but also increases chances of getting non-target loci (e.g pseudogenes, non-homologous locus). Our study shows that quality control steps are required to make sure the sequences included in the data sets are accurate. Indeed, clusters that contained the largest proportion of reads can contain frame shift mutations and/or stop codons or belong to non-target organisms (e.g. symbionts or parasites). Amplification of sequences from symbionts or parasites occurred almost exclusively when we used primers derived from the universal Folmer’ primers to amplify *COI* (LCO1490, HCO2198 ^29^) (up to 25.9% of the clusters obtained from positive PCR products and that passed the translation step). This shows that caution must be taken when using *COI* for a metabarcoding approach to assess species diversity.

The power of our approach coupled with its simplicity makes it attractive, but good practice designed to detect issues with Sanger sequences are still relevant. At least a translation to amino acids and a comparison to existing database (e.g. through BLAST) should be performed before sequence validation. While it has been underlined that pseudogenes, heteroplasmic sequences or sequences from symbionts or parasites may be obtained^4^, contamination during library preparation is less discussed. Indeed, contamination is difficult to detect and requires taxonomic knowledge and sequencing of both mtDNA and nuDNA markers to be distinguished from mtDNA introgression. Our results confirm that several markers should be sequenced for species diversity assessment to avoid underestimation of the number of species.

While it may be possible to identify cross-contamination or amplification of non-coding copies (e.g nuclear mitochondrial pseudogenes, numts), sorting out paralogs in which no stop codon or frameshift mutation is detected may be difficult. In this study, we found nine specimens for which we were not able to select just one from the two copies that passed our quality control steps. These specimens were thus represented by two sequences in the final MiSeq data set. Origin of these sequences is difficult to assess. PCR or sequencing mistakes cannot be ruled out and use of replicate sequencing may reduce the noise in data processing ^30^ but NUMTS or heteroplasms can also explain such pattern as NUMTS have already been identified in fig wasps ^31^. Differentiating heteroplasmic sequences from sequences of recent pseudogenisation is tricky and rely mostly on mitochondrial enrichment experiments ^32^. When compared to the total number of samples for which we managed to select only one cluster : 306 (*COI*), 261 (*Cytb*), 273 (*EF*) this result may appear negligible but this pattern could be problematic for groups in which coding NUMTS are frequent (e.g. grasshoppers ^12^ or longhorn beetles ^33^).With Sanger data sets, unrecognized co-amplification of heteroplasmic sequences or nuclear mitochondrial pseudogenes (NUMTS) have been frequently interpreted as the presence of cryptic species – especially in absence of taxonomic expertise - and have contributed to overestimating the number of unique species ^12^.

On the final trees (Sanger + Miseq sequences), 4.3% (*COI*) and 5.2% (*Cytb*) of the species for which at least one consensus sequence passed quality control were paraphyletic or clustered into two divergent groups of sequences (>7%). In some cases, for Cytb, such pattern could be explained by the fact that two primer pairs were used to generate the Sanger data set, while only one pair was used for the MiSeq part. However the same pattern was observed when the same pair was used for the Sanger and the MiSeq data set. All these cases are difficult to explain (PCR/sequencing errors, NUMTS, heteroplasmic sequences ?). They showed that filtering methods to select sequence clusters may influence the results and could lead to an overestimation of species diversity. In conclusion, these results are a reminder of how important it is to take a close look at the data, work in close relation with expert taxonomists and consider more than one marker for DNA-taxonomy or species diversity assessment.

To conclude, the approach presented here may contribute to the acceleration of the global efforts and may also contribute to improving the state of completeness and accuracy of the present database. We also advocate capitalizing on the huge investments made to construct barcode databases (BOLD) and in practice be conservative and pragmatic in maintaining the genes and the methods mastered by scientists, while shifting to next generation sequencing.

## METHODS

### Study group

We used the fig wasp genus *Ceratosolen* (Hymenoptera: Chalcidoidea: Agaonidae) as a test case because it is a relatively diverse genus of insects (encompassing an estimate of 230 species worldwide, of which only 71 are described). This genus pollinates *Ficus* species of the subgenus *Sycomorus*(158 described species worldwide) and is thoroughly studied by researchers working on figs. The genome of one species of *Ceratosolen* has been recently sequenced ^34^. Furthermore, in the last 20 years, we have developed a multigenic barcoding database using Sanger sequencing that encompass an unprecedented sampling of species. We have previously sequenced these specimens on *COI*, *Cytb* and *EF1*α.

### Sampling

One-hundred twelve species of *Ceratosolen* were included in the data set, of which about half (n=62) are undescribed. Three species were taken as outgroups : two in the genus *Kradibia* (sister group of the genus *Ceratosolen*^35^) and one in the genus *Tetrapus*. All material were collected alive from 1996 to 2015, fixed in 75% ethanol and identified morphologically by JYR.

*Sanger*: DNA from two to three specimens per species was extracted over the past 20 years.

*MiSeq* : On average, DNA from three specimens per species was extracted. A total of 369 individual specimens were included in the library.

### DNA extraction

DNA was isolated using the Qiagen DNeasy^®^ 96 Blood and Tissue Kit (Qiagen, Germany) according to manufacturer’s protocols. Individual specimens were incubated overnight at 56°C in the lysis buffer before performing the next extraction steps. In the end, DNA was recovered in a total of 100 µL of AE buffer (two elution steps of 50 µL AE buffer each).

With very few exceptions, sequences were obtained from the non-destructive extraction of a single wasp specimen (corpse kept as voucher). When destructive extraction was used, vouchers were selected among specimens sampled from the same tree and the same fig after careful identification by JYR. Destructive extraction was performed for the Miseq library. Vouchers are deposited at CBGP, Montferrier-sur-Lez, France.

#### Sanger data set

Two mitochondrial protein-coding genes [the 5’ end of the cytochrome c oxidase subunit I (*COI*) « barcode fragment » and part of the cytochrome b (*Cytb*)] and one nuclear protein-coding gene [elongation factor-1a (*EF1a*)] were included in the study. Amplification and sequencing protocols followed Cruaud et al. ^36^ for *Cytb* and *COI* and Cruaud et al. ^37^ for *EF1a*. The two strands for each overlapping fragment were assembled using Geneious v6.1.6 ^38^. All sequences that we obtained for the target species were included in the data set. Sequences were aligned using MAFFT v7.222 ^39^ (L-INSI option). Alignments were translated to amino acids using Geneious v6.1.6 to detect frameshift mutations and premature stop codons. Phylogenetic trees were inferred for each gene using RAxML v8.2.4 ^40^. Given that α and the proportion of invariable sites cannot be optimized independently from each other ^41^ and following Stamatakis’ personal recommendations (RAxML manual), a GTR + Γ model was applied to each gene region. We used a discrete gamma approximation ^42^ with four categories. GTRCAT approximation of models was used for ML boostrapping ^43^ (1000 replicates). Resulting trees were visualised and annotated using TreeGraph 2 ^44^. Following visual inspections of trees, contaminations (100% identical sequences for samples belonging to different species between which hybridization is not possible) were removed from the data set.

### Illumina MiSeq library preparation

Our library preparation approach involved two PCR steps with different primer pairs, as suggested in the Illumina protocol for 16S Metagenomic Sequencing Library Preparation. The first PCR step is performed to amplify the targeted DNA region. In this step, the primer pairs used contain a standard Illumina sequencing primer, a 0 to 3 bp “heterogeneity spacer” (as suggested in Fadrosh et al., ^45^) and the gene-specific primer (Fig. 1). The second PCR step is performed in order to multiplex individual specimens on the same Illumina MiSeq flowcell and to add necessary Illumina adapters. In this second step, primer pairs used contain the appropriate Illumina adapter allowing amplicons to bind to the flow cell, a 8-nt index sequence(as described in Kozich et al., ^46^) and the Illumina sequencing primer sequence. We used negative controls (DNA extraction and PCR) on each plate from the beginning to the end of sequencing.

*First PCR step* : each reaction contained 3 µL DNA template, 5 µL QIAGEN Multiplex PCR Master Mix (Qiagen, Germany) (including Taq polymerase, dNTPs and MgCl2), 0.5 µM forward primer, 0.5 µM reverse primer and 1 µL molecular biology grade water in a total volume of 10 µL. PCR conditions were 95°C for 15 min; 35 cycles of 94°C for 30 sec, 51°C for 90 sec and 72°C for 60 sec; and a final extension at 72°C for 10 min. All amplifications were completed on a Eppendorf Mastercycler ep gradient S thermocycler (Eppendorf, Germany). The same primers as for the Sanger data set were used. Two primer pairs for *COI* : LCO1490puc + HCO2198puc ^36^ (“*COI*-long”, Table 1) and UEA3 ^47^ + HCO2198 ^48^ (“*COI*-short”); one pair for *Cytb* : CB1 + CB2 ^49^ and one pair for *EF1a* : F2-557F + F2-1118R ^50^. Thus, four amplicons were generated per specimens. Amplicons were visualized on 1% agarose gels to quantify PCR success.

*Second PCR step* : During this step, amplicons were dual indexed with multiple identifiers (MIDs). Each pair of indices (i5 and i7) was unique to a PCR well, with the aim of assigning each sequence to a sample. PCR conditions were 95°C for 15 min; 10 cycles of 95°C for 40 sec, 55°C for 45 sec and 72°C for 60 sec; and a final extension at 72°C for 10 min.

Positive and negative PCR amplifications were pooled into tubes (1 tube per primer pair) and the resulting mixtures were subjected to gel electrophoresis on a 1.25% low-melting agarose gels. The bands corresponding to the PCR products were excised from the gel and purified with a PCR clean-up and gel extraction kit (Macherey-Nagel, Germany). Purified DNA was recovered in a total of 40 µL of NE buffer and quality and quantity of PCR fragments were determined by running 1 µL of each sample on a Agilent Bioanalyzer 2100 using the DNA 1000 LabChip kit (Agilent Technologies, USA). Each library (one per primer pair) was then quantified with the Kapa library Quantification kits (Kapa Biosystems, USA). The four librairies were then pooled equimolarly (0.06 nM of each gene region). The low concentration of the resulting library (0.24 nM) led to a high concentration of NaOH in the final solution after diluting with HT1. We therefore introduced 200 mM Tris-HCl pH7 to ensure that NaOH will be correctly hydrolyzed in the final solution. PhiX control library (Illumina) was combined with the amplicon library (expected at 5%) to artificially increase the genetic diversity and the library was paired-end sequenced on a MiSeq flowcell using a V3 MiSeq sequencing kit. Image analysis, base calling and data quality assessment were performed on the MiSeq instrument.

### Analyses of the MiSeq data set. Step 1, from read filtering to clustering (Fig. 2)

Quality control checks were performed on raw sequence data with FastQC v.0.11.2 ^51^. Overlapping paired-end reads were reassembled using FLASH v.1.2.11 ^52^ with default settings and extended maximum overlap length (300). When paired-end reads did not overlap (COI-long and EF1a, Table 1), forward and reverse reads were analysed separately. CUTADAPT v.1.2.1 ^53^ with default settings was used to sort paired reads by gene region and remove primers. *COI*-long and *EF1a* forward and reverse reads were quality trimmed (reads were truncated at the first position having quality score < 21) using VSEARCH v.1.8.1 (available at https://github.com/torognes/vsearch). After removing primers and quality filtering, fastq files were converted to fasta files and sequences less than 150 bp in length were filtered out using VSEARCH. Remaining sequences were dereplicated and putative chimeric sequences were removed using VSEARCH. *Cytb* and *COI*-short sequences were then trimmed by length using PRINSEQ v.0.20.4 ^54^ (minimum length = 400 bp; maximum length = 550 bp). Illumina sequences were then clustered using SWARM v.2.1.6 ^55^ and CAP3 ^56^ with default settings. Finally, clusters containing less than 10 sequences were excluded from the data sets using VSEARCH. Difference in read lengths due to quality trimming leaded to an overestimation of the number of clusters by SWARM for *COI*-long and *EF1a* forward and reverse reads. Therefore only the results obtained with CAP3 were subsequently analysed.

### Analyses of the MiSeq data set. Step 2, quality control of clusters of reads (Fig. 3)

For each gene region, the consensus sequence of each cluster was aligned with the corresponding Sanger data set using MAFFT v7.222(default parameter). When paired-end reads did not overlap (*COI*-long and *EF1a*), clusters of reads 1 and clusters of reads 2 were analysed separately. At this step of the process, 6 data sets were assembled : *COI* Sanger + *COI*-short MiSeq, *COI* Sanger + *COI*-long MiSeq reads 1, *COI* Sanger + *COI*-long MiSeq reads 2, *Cytb* Sanger + *Cytb* MiSeq, *EF* Sanger + *EF* reads 1 MiSeq, *EF* Sanger + *EF* reads 2 MiSeq.Alignments were translated to amino acids using Geneious v6.1.6 to detect frameshift mutations and premature stop codons. Non-coding sequences were removed from the data set. NCBI-BLAST was used to identify sequences that did not belong to the target group. Phylogenetic trees were then inferred for each gene region using RAxML. Resulting trees were visualised and annotated using TreeGraph 2 ^44^. Visual inspection of trees was carried out to identify contaminations, which were subsequently removed from the data sets. Reads 1 and 2 for COI and EF were then merged into a single data set (gaps were inserted between the non overlapping reads). Finally,*COI*-long and *COI*-short data sets were merged and potential discrepancies were pointed out. MEGA7^57^ was used to calculate average divergence (p-distance) between sequence groups for problematic cases.

## ACKNOWLEDGEMENTS

We thank Maxime Galan (INRA, CBGP, France) and Adrien Vigneron (Newcastle University, UK) for technical advises; Laure Benoit (CIRAD, CBGP, France) and Guenaëlle Genson (INRA, CBGP, France) for their help with lab work; Hélène Vignes (CIRAD, GPTR, France) for sequencing of the library as well as Alexandre Dehne-Garcia (INRA, CBGP, France) for his help with the CBGP HPC computational platform on which analyses were performed. We thank Anthony Bain (Taipei University), Yang Da-Rong, Zuodong Wei and Peng Yan-Qiong (Xishuangbanna Botanical Garden, China), Rhett Harrison (CTFS, Malaysia), Rong Chien Lin (University Taipei, Taiwan), Emmanuelle Jousselin, Carole Kerdelhué (CBGP Montpellier, France), Marc Pignal (MNHN Paris, France), Etienne Randrianjohany (CNRE, Madagascar),Simon van Noort (Iziko Museums of South Africa), Rosichon Ubaidillah (LIPI Bogor, Indonesia) for providing samples. This work was supported by the network Bibliothèque du Vivant funded by the CNRS, the Muséum National d'Histoire Naturelle and the Institut National de la Recherche Agronomique.

## AUTHOR CONTRIBUTIONS

JYR and AC designed the study. JYR, LJR and AC collected the samples. PC performed laboratory work and designed the custom workflow. PC, JYR and AC analysed the data.. PC, JYR, LJR and AC wrote the manuscript.

## ADDITIONAL INFORMATION

The authors declare no competing financial interests.

